# Network controllability: viruses are driver agents in dynamic molecular systems

**DOI:** 10.1101/311746

**Authors:** Vandana Ravindran, Jose Carlos Nacher, Tatsuya Akutsu, Masayuki Ishitsuka, Adrian Osadcenco, V Sunitha, Ganesh Bagler, Jean-Marc Schwartz, David L Robertson

## Abstract

In recent years control theory has been applied to biological systems with the aim of identifying the minimum set of molecular interactions that can drive the network to a required state. However in an intra-cellular network it is unclear what ‘control’ means. To address this limitation we use viral infection, specifically HIV-1 and HCV, as a paradigm to model control of an infected cell. Using a large human signalling network comprised of over 6000 human proteins and more than 34000 directed interactions, we compared two dynamic states: normal/uninfected and infected. Our network controllability analysis demonstrates how a virus efficiently brings the dynamic host system into its control by mostly targeting existing critical control nodes, requiring fewer nodes than in the uninfected network. The driver nodes used by the virus are distributed throughout the pathways in specific locations enabling effective control of the cell via the high ‘control centrality’ of the viral and targeted host nodes. Furthermore, this viral infection of the human system permits discrimination between available network-control models, and demonstrates the minimum-dominating set (MDS) method better accounts for how biological information and signals are transferred than the maximum matching (MM) method as it identified most of the HIV-1 proteins as critical driver nodes and goes beyond identifying receptors as the only critical driver nodes. This is because MDS, unlike MM, accounts for the inherent non-linearity of signalling pathways. Our results demonstrate control-theory gives a more complete and dynamic understanding of the viral hijack mechanism when compared with previous analyses limited to static single-state networks.

## Introduction

To replicate viruses are entirely dependent on the host system they infect. This involves a high degree of virus-host specificity at the molecular level. For example, recognition of receptors on specific cell types by virus molecules is key to cell-entry1, and targeting of host molecules is necessary to exploit intra-cellular ‘machinery’ to replicate. To achieve this, virus molecules must interact with many host molecules through a complex network of mostly PPIs^2^. In turn the host response to infection involves anti-viral factors, and subsequent virus response to host response, leading to a complex entanglement of virus and host interactions^3^.

With the availability of virus-host interaction data sets, such as VirHostNet^4^, HHID^5^ and HCVpro^6^, it is now possible to study viral infection as networks. For instances, human-pathogens protein-protein interactions (PPI) networks have provided a global view of strategies used by different pathogens to subvert human cellular processes and infect them^7,8^, and HIV-human interaction networks have been investigated to provide insights about host-cell subsystems that are perturbed during infection and investigate approved drug targets^9–11^. It is clear that the specific intra-cellular functions required by a virus to replicate and maintain an infection are the important focus for the virus, and the individual molecular interactions with the host system are just a means to this end^12^.

Linked to networks, graph theory is widely used as a model to describe and visualize perturbation in host cellular systems at the molecular level. For example, the tendency of ‘hubs’ (highly connecting proteins) and high centrality proteins, to be targeted by viruses has been highlighted^7,13,14^. However, in some cases these can be explained by the over-representation of highly-connected molecules in the host function being targeted^9^. Unrealistically the majority of these studies analyse and visualise the virus-host relationships in static networks, i.e., all of the interactions represented as active, disregarding the temporal and spatial nature of infection. In reality the virus must interact with a complex non-linear dynamic host system. Study of the dynamic nature of infection tends to be limited to specific sub-cellular systems, for example, using logical models in the context of T-cell signalling^15^.

Dynamic systems control theory has emerged as a mathematical framework for understanding how best to control an engineered system. Control theory has largely been applied to study complex networks and identify ways to control network behaviour^16–20^. The aim is to identify the minimum number of inputs, termed ‘driver’ nodes, that can push the system from any initial state to any final state in finite time. Past studies have employed control theory to identify the driver proteins significantly enriched in human diseases like cancer^21–25^ and other biological functionalities^26–29^.

Viral infection is unique as a system for the study of the applicability of controllability to a natural system, as the virus makes many interactions with the host system and is explicitly controlling host functions. Past studies have explored the use of control theory in biology. However, in these systems only a few molecules are involved in any instance, e.g., in the case of disease or drug therapy based studies, and so studying all known disease-associated molecules or all drug-targets at the same time has little biological meaning^22,30^. In studies where viruses have been investigated explicitly the virus-host interactions have not been included in the network^26,27^, and so control theory has been partially applied to the system, and the effect of inclusion of viral proteins has not been modelled.

In this work, we have modelled HIV-1 and HCV host interactions as dynamic networks both with and without the virus-host interactions included. We examine these networks from a controllability perspective and test whether viruses follow the principles of control theory during hijack of a host system (1). This study of viral infection from a controllability perspective permits, for the first time, testing of the applicability of this mathematical framework to intra-cellular networks and discrimination of available control models (maximum matching versus minimum dominating set^17,18^) using a dynamic network, consisting of two states: normal/uninfected and infected. Our results lead to novel understanding of the infection mechanism limited to single-state networks, demonstrating the applicability of control theory to the study of infection and validating its use in the study of intra-cellular networks.

## Results

### Identification of driver nodes in the uninfected network

We first identified the driver nodes without the presence of virus in a large directed signalling network comprised of 6339 nodes and 34813 interactions, data from Vinayagam *et al*.^26^. In this network the nodes represent proteins and the edges/links represent directed signal flow between them (Supplementary Data file 1). The minimum number of driver nodes (*N_D_*) were identified and compared using two established models (i) minimum dominating set (MDS)^18^ and (ii) maximum matching (MM)^17^ (2). This analysis classified 1398 (22%) of the nodes as driver nodes based on the MDS method, compared to 2282 (36%) of them classified by MM (Table 1). This captures the ‘ease’ with which the network can be controlled and demonstrates that MDS yielded many fewer driver nodes compared to MM.

**Table 1.** Classification of minimum dominating set (MDS) and maximum matching (MM) driver nodes.

|  | MDS model |  |  | MM model |  |  |
| --- | --- | --- | --- | --- | --- | --- |
|  | Normal | Infected |  | Normal | Infected |  |
|  |  | HIV | Human |  | HIV | Human |
| **Driver nodes** | 1398 | 1232 |  | 2282 | 2264 |  |
| **Critical** | 874 | 12 | 688 | 377 | 1 | 266 |
| **Intermittent** | 1250 | 5 | 1231 | 3330 | 2 | 3443 |
| **Redundant** | 4215 | 5 | 4420 | 2632 | 19 | 2630 |

Since the minimum driver node set obtained by both these models is not unique, we determined the role of a node as a driver node and classified each node as ‘critical’ if it was present in all driver node sets, ‘intermittent’ if it was present in some driver node set and ‘redundant’ if the node was never part of any driver node set. We find 14% of the nodes were critical in MDS, while only 6% of the nodes were classified as critical using MM model. Similarly, the MDS classified more nodes as redundant compared to MM. On the other hand, the MM model classified most of the nodes as intermittent (53%) compared to MDS (20%) (Table 1, Figure 1). Collectively this explains why fewer overall driver nodes are required to control the network with MDS. MDS, thus, better reflects optimal control of the signalling network by identifying fewer driver nodes compared to MM.

**Figure 1.**
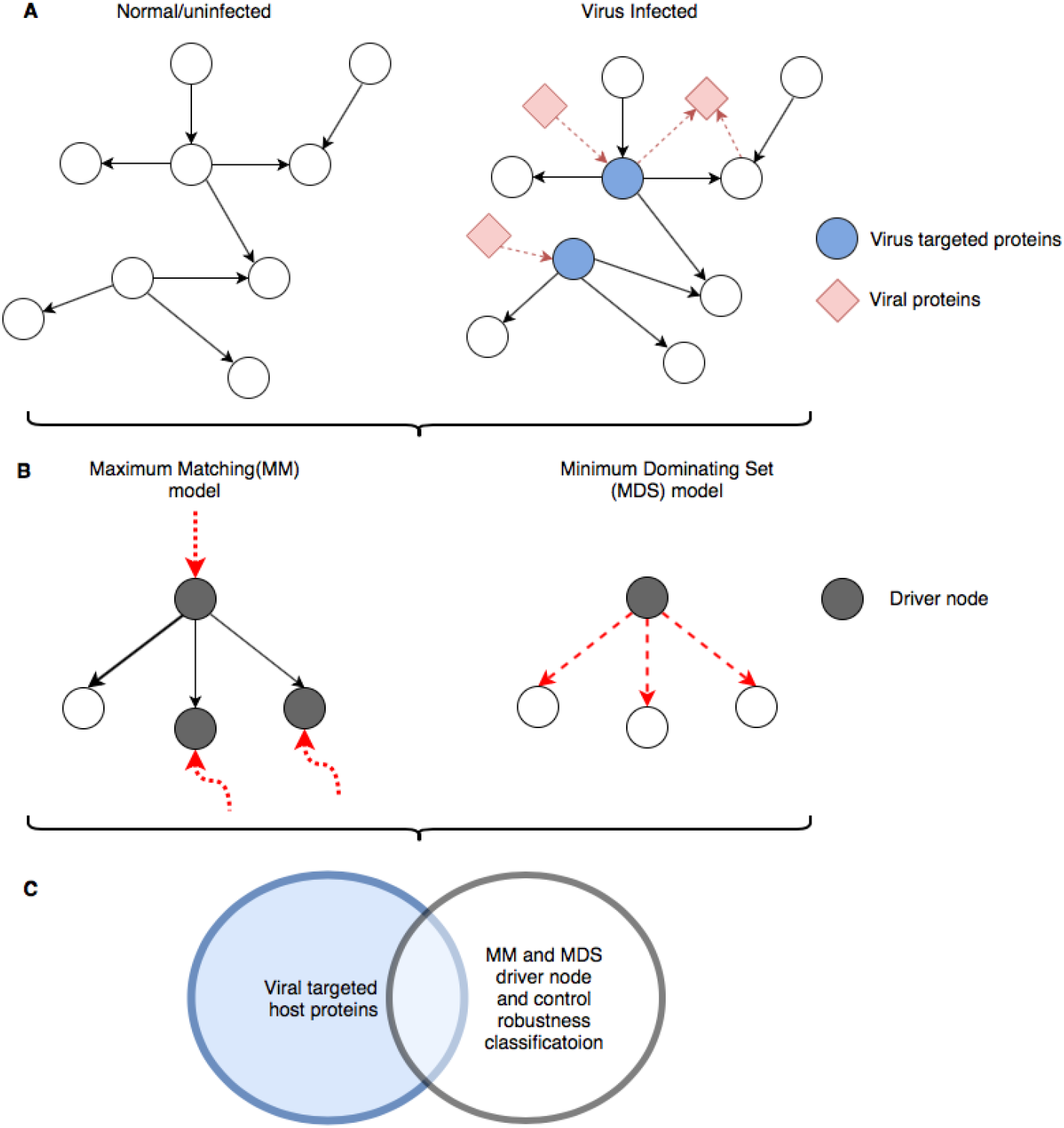
Schematic representation of our network controllability analysis for virus infection. (A) Example of normal and infected network. (B) Driver node identification using maximum matching (MM) and minimum dominating set (MDS) models. The red dotted arrows indicate inputs to the driver nodes. The bold arrow indicates the matched edge on the maximum matching. (C) Comparison of host proteins virus interacts with and the identified driver nodes.

**Figure 2.**
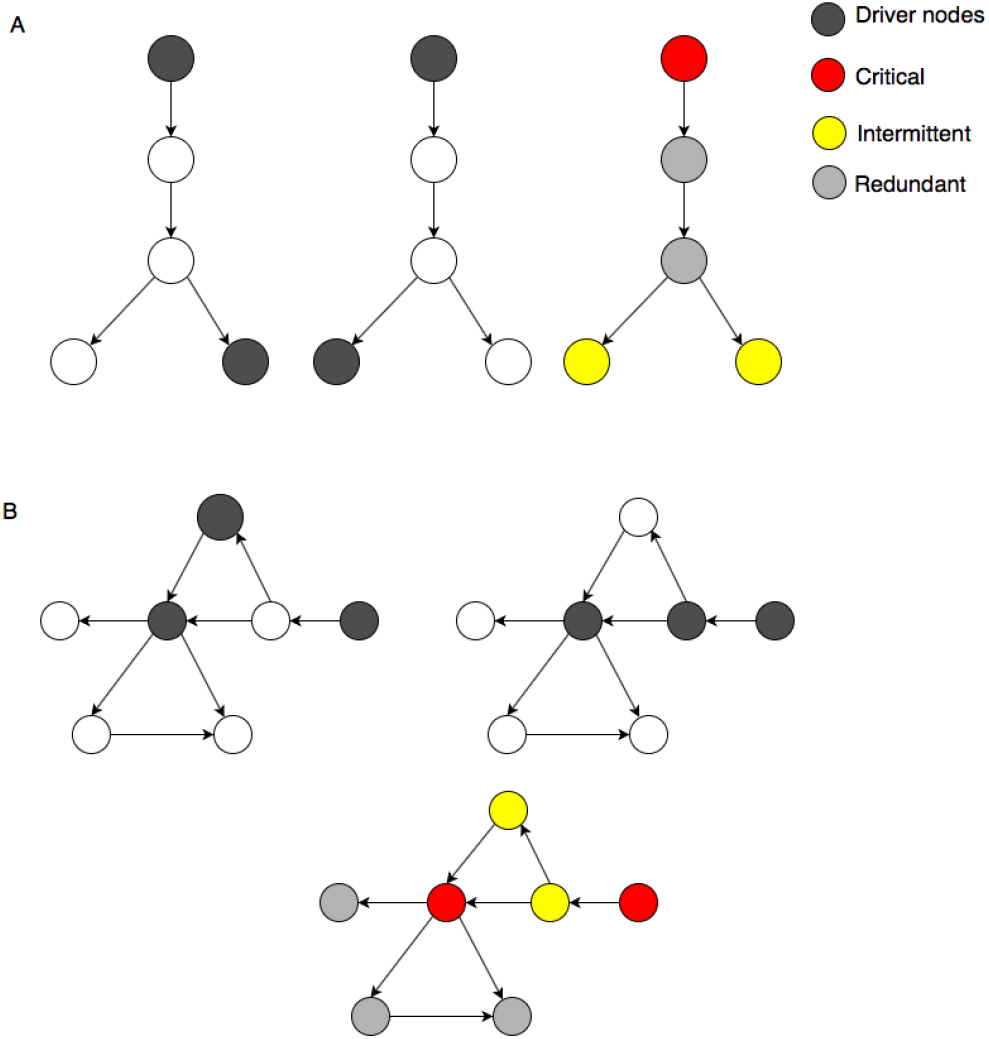
Example of identification and classification of driver node sets into whether a node is always present in these sets (critical driver node), occasionally present (intermittent driver node) or never a driver node (redundant) for the (A) maximum matching (MM) and (B) minimum dominating set (MDS) models. See key for node designations.

The driver nodes identified by MM tend to have a zero in-degree, corresponding to receptors on the cell that extracellular molecules (ligands) interact with in order to convey signals to the cell via the signalling pathways. MM only identifies receptors as the driver nodes because this model of control tends to prioritize as drivers those nodes at the beginning of long linear chains, which are controlled externally. The internal nodes of these chains are controlled internally through maximum matched links, whose coupling are consistent with linear response systems as described in the linear mathematical equations shown in Liu *et. al* (2011)^17^. Molecular signalling systems, however, are inherently non-linear. Thus, the MM will only detect the initial nodes, such as receptors, as the driver nodes as observed in a few signalling studies^26,30^. On the contrary the MDS method does not require structural controllability, here a single integrator node that receives a unique signal from an input link makes the node controllable. Given that the dynamics underlying biological networks is non-linear and that the MDS better reflected the biological reality by identifying few driver nodes, it is thus a more appropriate model for controllability of an intra-cellular network.

In order to assess the roles of proteins in the context of cell signalling, we classified them as either signalling proteins, kinases, receptors and transcription factors^26^. In total the proteins were classified into 1006 signalling proteins, 545 receptors, 366 kinases and 1150 transcription factors (Supplementary Data file 2). We observed that the critical driver nodes obtained through the MDS model played diverse roles in signalling processes. They were highly enriched as receptors, consistent with MM (see Supplementary Material, Tables S1 and S2). MDS also showed enrichment for signalling proteins and kinases. The redundant nodes were enriched for transcription factors while the intermittent nodes showed no distinct enrichment.

### HIV-1 targeted driver nodes in the uninfected network

Next we looked at the association of virus proteins with driver nodes to identify if they are preferentially targeted by the virus. Out of 6339 proteins in this human PPI network, 2529 nodes have been reported to be targeted by HIV-1. Of the different MDS driver node classes targeted by HIV-1 we observed that, compared to random samples, 50% of the critical driver nodes were significantly targeted by HIV-1 (Z-score= 6.50) (Table 2), rather than intermittent and redundant nodes, indicating that critical driver nodes are preferentially targeted by the virus.

**Table 2.** HIV-1 targets among MDS driver nodes. Numbers of observed critical, intermittent or redundant nodes were compared to 1000 random samples.

| Node type | Observed | Percentage | Random mean | Z-score | P-value |
| --- | --- | --- | --- | --- | --- |
| **Critical** | 438 | 50.11 | 349.38 | 6.5 | 8.03E-011 |
| **Intermittent** | 489 | 39.12 | 498.4 | -0.61 | 0.542 |
| **Redundant** | 1602 | 38.01 | 1681.45 | -4.41 | 1.03E-005 |

To gain more insights into the role of driver nodes and their preference as viral targets, we propose a novel metric for MDS model: the control centrality (CC) metric that measures the ‘power’ of a node in controlling other nodes or the number of controllable nodes in the network. For an MDS, control centrality of a node v is *k_out_* + 1, in which *k_out_* denotes the node out-degree. We used this method to compute the control centrality metric for critical, intermittent and redundant nodes (Table 3). For comparison, we averaged this measure and observed that the control centrality of critical driver nodes was more than that of intermittent and redundant nodes. Given our observation that the critical driver nodes were predominantly receptors (45%) and thus of major importance, i.e., ‘powerful’ in controlling the system based on CC, this explains their preference as targets by HIV-1.

**Table 3.**
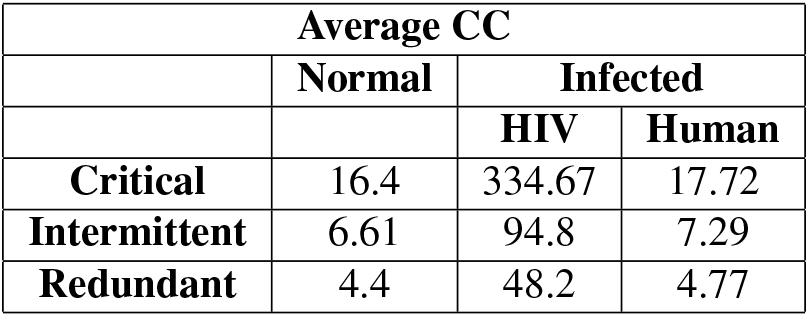
Control Centrality (CC) metric of MDS driver nodes.

### Identification of driver nodes in the HIV-1 infected network

Driver nodes were again identified based on the MDS and MM models this time with the inclusion of the virus-host interactions in the directed network (Supplementary Data file 1). For MDS, 1232 (19%) of the nodes were driver nodes, while for MM 2264 (36%) of the nodes were driver nodes (Table 1), results that are approximately comparable to the uninfected network. MDS identified fewer driver nodes compared to MM in the infected networks as well. In the MDS model, 166 nodes lost their driver node status upon infection with HIV, compared to a decrease of only 18 driver nodes for the MM model (Table 1, Supplementary Data file 2)

While there is an overall decrease in the number of critical driver nodes identified by both the models, intermittent and redundant nodes exhibited almost no change for the uninfected and infected states (Table 1). Importantly, it is only for MDS that the majority of HIV-1 nodes are critical driver nodes (12 of 22), while for MM only 1 of 22 have this property (Table 1). These ‘main’ 12 HIV-1 proteins, not unexpectedly, are: Nef, gp120, Tat, Pr55(gag), caspid, Rev, gp41, Vpr, Vif, retropepsin, Vpu and p51. In addition the critical nodes identified by MM tend to be more ‘peripheral’ in the network as they correspond to the zero in-degree nodes (Figure 3). Collectively these results confirm that MDS is matching the viral control of the system much more accurately than MM.

**Figure 3.**
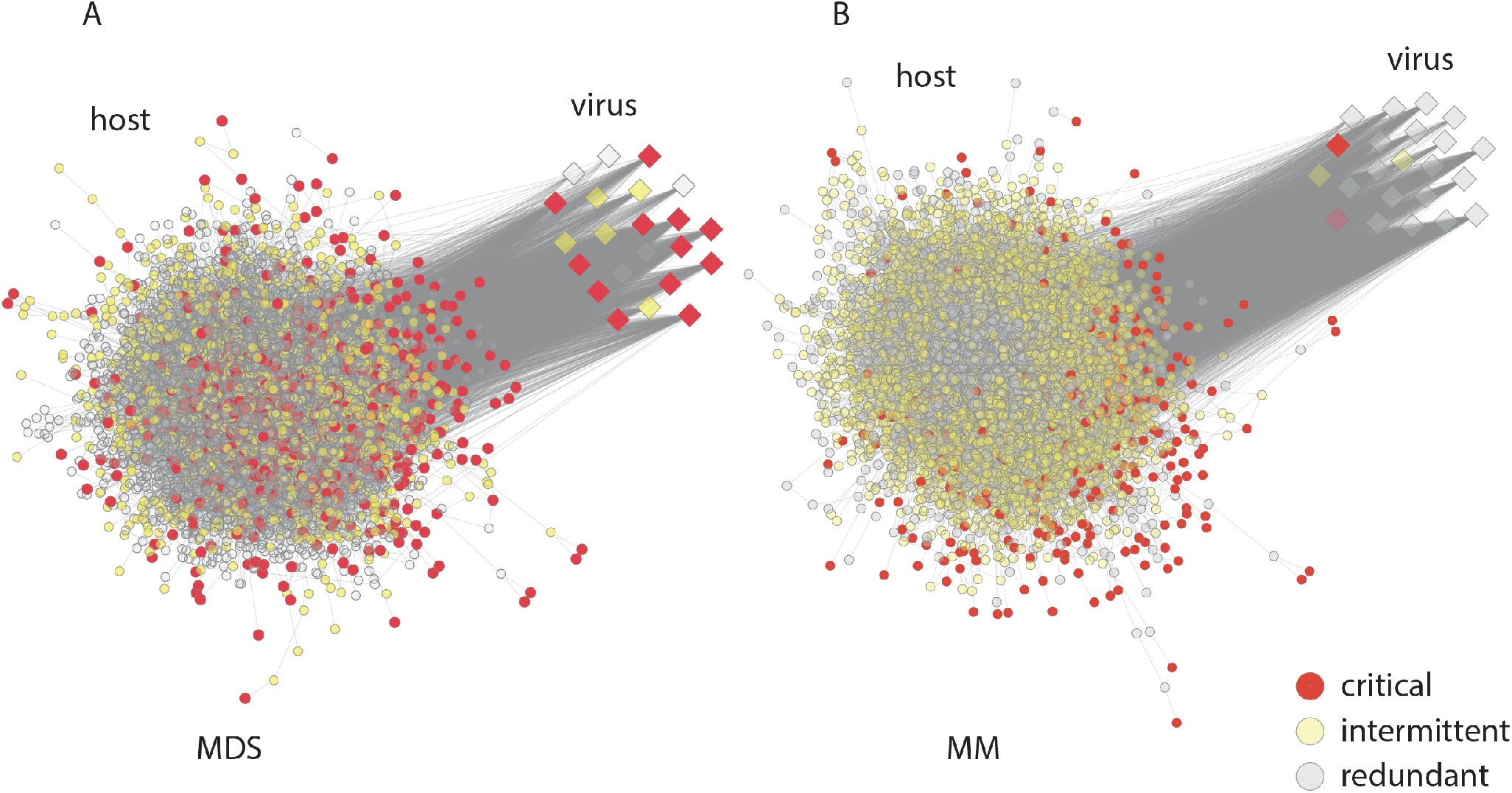
Visualisation of critical, intermittent and redundant driver nodes (see key for designations) for the signalling network infected with HIV-1 for (A) the minimum dominating set (MDS) and (B) maximum matching (MM) models. Host and viral proteins are shown as circles and squares respectively. Grey lines denote an interaction.

These results indicate that inclusion of the virus has increased the controllability of the network, i.e., the virus set of interactions increases the total number of interactions facilitating the ‘hijacking’ of cell, and controlling the network more efficiently. Further, the control centrality analysis for the HIV-1 infected network indicates that HIV-1 molecules are an order of magnitude more ‘powerful’ than host molecules (Table 3).

In the HIV-1 infected network, the MDS model identified 42 nodes as new critical control nodes, while 646 nodes were ‘preserved’ critical nodes, i.e., they retained their driver node status even after HIV-1 infection (Supplementary Material, Figure S1(a), and Supplementary Data file 3). Again, this indicates that the controllability model is capturing the biological signal in terms of the virus mechanism of control.

We also looked at those critical driver nodes that are HIV-1 targets. 438 critical driver nodes were HIV-1 targets in the uninfected network, which reduced to 271 in the infected network (Supplementary Material, Figure S1(b)). This change is presumably due to the inclusion of the HIV-1 host interactions causing many host molecules to lose their critical driver node status. This indicates the virus is fundamentally changing the controllability of the host system. Interestingly, after infection 248 HIV-1 target proteins still preserved their critical driver node status (Supplementary Material, Figure S1(b) and Supplementary Data file 3). These 248 critical driver nodes are not just important from control point of view but are also the nodes that are HIV-1 targets and could act as potential drug/intervention targets.

We investigated the biological properties of the preserved critical driver node targets by submitting them to the Reactome Pathway database^31^ for enrichment analysis. These critical nodes are enriched for 1341 pathways in the Reactome database. As expected, given the nature of the data, the vast majority were active in signal transduction followed by the immune system. Interestingly, the critical driver nodes seem to play roles in many other cellular subsystems, ranging from cell cycle to developmental biology, disease, programmed cell death and matrix organisation among others, though not as pronounced as in the first two (Table 4). Using the preserved critical proteins, the enriched pathways were analysed to ascertain the top 10 for each set of proteins (Table 4). The list was curated using the FDR rate, p-value for significance, ratio of hit proteins in pathway and similarity percentage between proteins within the enriched systems. Not surprisingly given HIV-1’s life cycle, the top five pathways are specific to the immune system, though most of them also have moderate overlap with RAF/MAP kinase cascades in signal transduction. The next three pathways exclusively deal with signal transduction, followed by programmed cell death, and the last pathway deals with gene expression, transcription in particular. These analyses ascertain the different pathways that the preserved critical driver nodes are involved in and demonstrate the MDS model is a useful method to identify different intra-cellular pathways that HIV-1 interacts with.

**Table 4.** Top 10 Reactome enriched pathways for MDS preserved critical driver proteins between the normal/uninfected and HIV-1 infected network.

| Pathway name | Total proteins in pathway | Matching proteins | p-value | FDR |
| --- | --- | --- | --- | --- |
| Toll-Like Receptor Cascades | 141 | 14 | 9.27E-06 | 1.23E-04 |
| Signalling by Interleukins | 460 | 43 | 1.15E-14 | 2.21E-12 |
| CD28 co-simulation | 29 | 6 | 8.01E-05 | 5.61E-04 |
| Fc epsilon receptor (FCERI) signalling | 405 | 32 | 2.96E-09 | 1.75E-07 |
| Signalling by B Cell Receptor (BCR) | 270 | 23 | 1.68E-07 | 5.89E-06 |
| Signalling by NGF | 421 | 45 | 1.11E-16 | 5.30E-14 |
| Signalling by PDGF | 328 | 33 | 3.32E-12 | 3.52E-10 |
| Signalling by Wnt | 230 | 20 | 8.29E-07 | 2.24E-05 |
| Intrinsic Pathways for Apoptosis | 41 | 10 | 7.61E-08 | 2.89E-06 |
| SMAD heterotrimer regulates transcription | 32 | 7 | 1.43E-05 | 1.57E-04 |

### Control robustness: before and after infection

To analyse the robustness to control of the signalling network we classified each node into one of the following three categories26: (1) ‘indispensable’, i.e., positive control factor, if we have to control more driver nodes in its absence; (2) ‘dispensable’, i.e., negative control factor, if we have to control fewer driver nodes in its absence; and (3) ‘neutral’ control factor if in its absence there is no change in the number of driver nodes to be controlled (Supplementary Data file 2).

Nodes were categorised and compared based on the dispensable classification in both networks using MDS and MM, and the minimum number of the driver nodes (*N_D_*) was calculated. Interestingly, MDS classified a much smaller number of nodes as indispensable (503) or dispensable (770) compared to MM (1330 versus 2347 respectively), with twice the number of neutral nodes identified by MDS compared to MM (Table 5). As a consequence MDS performs much more efficiently than MM. A similar trend was seen in the HIV-1 infected network with a smaller number of nodes classified as indispensable (397 human and 11 HIV) or dispensable (719 human and 3 HIV-1) compared to MM (1331 human and 19 HIV-1 proteins verses 2346 human and 1 HIV-1). In addition, on comparing the difference in node characterisation in both states (normal/uninfected and infected), the indispensable nodes reduced by about 20% to 397 for the MDS model but showed no change for the MM model (Table 5). Again these results indicate MDS is a more appropriate model of control for the infected network than MM.

**Table 5.**
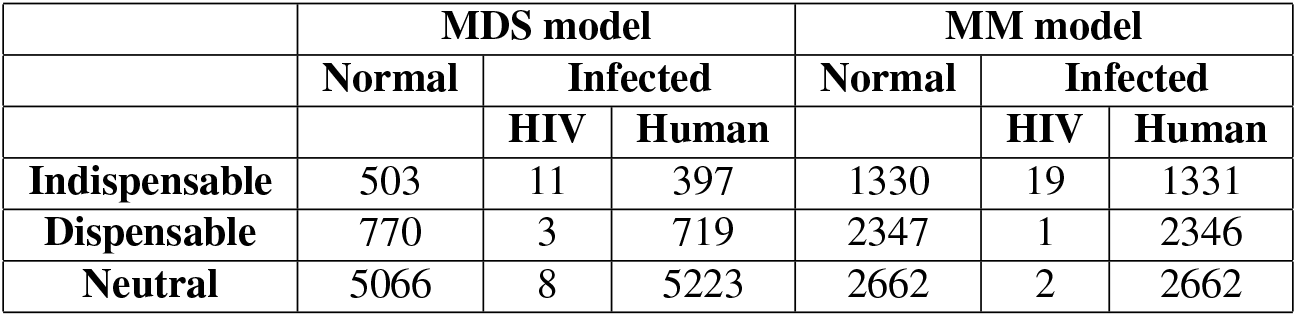
Control robustness analysis: Classification of nodes based on deletion studies between normal/uninfected and HIV-1 infected networks.

Comparing the alternative characterisation of nodes obtained from the MDS model for identifying the overall node control profile, we find that all of the indispensable nodes were critical and all redundant nodes were neutral in both networks; and have deduced a mathematical proof for this property of MDS (See Methods and Supplementary Note). From a control point of view, indispensable nodes not only determine the ease of control, but also are the driver nodes, since the ability to efficiently control a system is determined by the minimum number of driver nodes. Further, from a biological perspective, these nodes were frequently targeted by HIV-1 (62%) compared to other node classes (see Supplementary Material, Table S3).

We also performed controllability analysis using the MDS model for an HCV infected network to validate our findings with another virus (Supplementary Note).

Between uninfected and HCV-infected networks, a higher number of proteins (836 nodes) share critical driver node status, most likely due to the fact that the HCV viral targets fewer proteins (given available data). 16 proteins become critical after HCV infection, while 38 proteins lose their critical node status (Supplementary text-1 Figure S2). We analysed 65 preserved critical driver nodes for pathway enrichment (Supplementary text-1 Table S5). Many of the top 10 pathways were similar to that obtained for HIV-1. Particularly, CD28 co-stimulation, PDGF signalling, signalling by NGF, Fc epsilon receptor signalling and Apoptosis pathway. All of the immune system pathways and apoptosis pathways are shared amongst both viruses. Regarding signal transduction pathways, the viruses have only NGF and PDGF signalling in common, whilst the remaining signalling pathways seem to be particular to HCV only (Supplementary text-1, Figure S4). Interestingly about 40 proteins are common targets among HIV-1 and HCV (Supplementary text-1, Table S6).

## Conclusion

We have compared two network states (uninfected and infected) from a controllability perspective and find that the MDS model more effectively captures the dynamics of viral infection than MM. The performance of MDS validates, for the first time, the applicability of the control theory framework for the study of intra-cellular signalling networks and as a model for studying viral use of host cells. Our results clearly demonstrate the way in which a natural control system (virus exploitation of a host cell) can be used to understand the control of information flow in intra-cellular networks. This hints at the possibility of learning how to synthetically control complex biological systems. In terms of understanding infection, the virus is mainly ‘driving’ the network by exploiting it’s usual dynamics, i.e., tending to target the existing critical driver nodes. Interestingly, we demonstrate that indispensable nodes, the positive control factors, are always the MDS critical driver nodes. With the addition of the high-powered viral proteins (as measured by our CC analysis), this control is achieved more efficiently with fewer host molecules acting as critical driver nodes. The MDS proteins, specifically the critical driver nodes, are effectively ‘central’ molecules in terms of information flow in the system^32^ being enriched significantly for proteins that are highly-connected and often multi-functional. As the virus is mostly interacting with the host systems to replicate itself, involving the up and down regulating of specific host functions, control theory offers an ideal model for the study of this control of information flow, and we believe opens a new discipline of viral control theory. This has the potential to enhance our ability to interfere with infection, for example, by better understanding of the aberrant functions stimulated as a result of infection will be helpful in terms of treating the side-effects/symptoms of infection. Fully understanding the entanglement of viruses with host systems will be the key to limiting their harmful tendencies.

## Methods

### Data procurement and network construction

The human directed signalling network was obtained from Vinayagam *et al*.^26^. It consists of 6,339 proteins and 34,813 interactions. Interaction direction represents potential signal flow between interacting proteins, which was predicted using a Naïve Bayesian Classifier. The classifier assigns a score to each interaction ranging from 0.5 to 1 if there is signal flow, otherwise it assigns a score of 0^33^ (Supplementary Data file 1).

### HIV-1 infected network

HIV-1 interactions were obtained from HHID^5^. A total of 15,230 interactions were retrieved and were further curated by ignoring the number of publications, counting each reaction type only once and selecting only those interactions that had shared nodes with the signalling network. In this human network 2,529 proteins interact with HIV, resulting in a total of 5,811 additional interactions. The directions of the HIV-host interactions were assigned using the method provided by MacPherson *et al*. (2010)^12^, where each HHID interaction was assigned a direction based on its interaction type. Direction represents whether the virus protein acts upon the host or vice versa. For example, “Nef activates ACHE” would be given a forward direction as the virus protein acts upon the host, whereas “Nef is activated by ACHE” would be attributed a backward direction, since it is the host protein that activates the virus protein^12^. The final list of HIV-human PPIs was formed of 6,361 proteins and 40,625 interactions (Supplementary Data file 1). See Supplementary Material file for details on the HCV data.

### Driver node identification: minimum dominating set method

Driver nodes are identified by calculating the minimum dominating set for a given network. For a graph *G*(*V, E*), where *V* is set of nodes and *E* is set of edges, a subset *S* ⊆ *V* is called dominating set (DS) if every node in *V* is either an element of *S* or is adjacent to an element of *S*. That is for a directed graph, any node *v* ∈ *V, v* ∈ *S* holds or there is a node *u* ∈ *S* such that there exists a directed edge (*u, v*) ∈ *E* then we say that *v* is dominated by *u*. Then *S* is dominating set if each node in *V* is either in *S* or dominated by some node in *S*. A minimum dominating set (MDS) is a dominating set with the minimum number of nodes. The MDS forms the driver node set. Since the computation of MDS is NP-hard, we used integer linear programming (ILP) to compute the MDS by assigning 0 – 1 variable to each vertex, where 1 is if *v* is part of MDS else 0^18^. If a node is always present in all MDS, it is a critical driver node, occasionally present in MDS then it is an ordinary driver node and if a node is never part of any MDS then it is a redundant/non-driver node. To address critical controllability in large-scale PPI networks analysed in this study, we used a fast algorithm adapted to directed networks that uses efficient graph reduction using heuristics and mathematical propositions^34,35^. The algorithm for the undirected case was used to analyse large protein interaction networks integrating transcriptome^28^.

### Driver node identification: maximum matching method

Driver nodes were identified using the controllability package by Liu. *et al 2011*^17^. The algorithm converts the network into a bipartite graph and identifies the maximum matching for a digraph using Hopcroft-Karp algorithm. The unmatched nodes are the driver nodes and the minimum number of driver nodes is denoted by *N_D_*. Thus, if a node is always unmatched then it is a critical driver node (CDN), if it sometimes matched and unmatched it is an ordinary driver nodes (ODN) while if it is always matched it is a redundant/non-driver node (NDN).

### Calculation of control centrality measure of nodes in minimum dominating set

Control centrality is a measure that determines the power of a node to control its sub-systems or other nodes. Mathematically, for a directed network the control centrality of a node v is *k_out_* + 1. The reasoning for this count is an MDS driver node controls itself and the outgoing edges independently^18,34^.

### Classification of nodes based on their impact to size of driver nodes set in MDS

The nodes in the network were classified based on their impact on size of drive node set *N_D_* or dominating set (DS). For a network *N* a node was deleted at a time and the MDS was computed for the new network *N*′. If 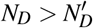 then the node is indispensable. If 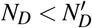 then dispensable and if 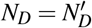 the node is neutral.

We also have shown that in an MDS, every indispensable node is critical and every redundant node is neutral (Supplementary Note). These properties are different from those obtained by the MM method^26^.

**Proposition 1.** *In MDS, every indispensable node is critical*.

*Proof*. We prove the proposition by contraposition. That is, we show that if *v* is not critical, *v* is not indispensable. Let *G*(*V, E*) be an undirected graph. Suppose that *v* ∈ *V*is not a critical node in *G*(*V, E*). Then there must exist an MDS *U* ⊂ *V* such that *v* ∉ *U*. Let *G*′(*V*′, *E*′) be the undirected graph obtained by removing v (i.e., deleting v and its connecting edges). Then *U* is a dominating set of *G*′(*V*′, *E*′) because each node *u* ∈ *V* – *v* was dominated by some node *w* ∈ *U* (*w* ≠ *v*) in *G*(*V, E*). It means that the MDS size of *G*′(*V*′, *E*′) is not larger than that of *G*(*V, E*). Therefore, *v* is not indispensable.

**Proposition 2.** *In MDS, every redundant node is neutral*.

*Proof*. Let *G*(*V, E*) be an undirected graph. First we show that any redundant node is not indispensable. Suppose that *v* is a redundant node in *G*, which means that *v* does not appear in any MDS of *G*. Let *G*′ be the graph obtained by removing *v* from *G*. Let *U* ⊂ *V* be an MDS of *G*. Then *v* ∉ *U* holds, which implies that *U* remains to be a dominating set in *G*′. Therefore, the size of an MDS does increase after removal of *v* and thus *v* is not indispensable. Next we show by contraposition that any redundant node is not dispensable. Let *W* be an MDS of *G*. Suppose that *v* is dispensable. Let *G*′ be the graph obtained by removing *v* from *G*. Since *v* is dispensable, there must exist an MDS *U* ⊆ *V* – {*v*} of *G*′ such that |*U*| = |*W*| – 1. Let *U*′ = *U*∪{*v*}. Clearly, *U*′ is an MDS of *G*, which implies that *v* is not redundant. By combining the above two properties, we can see that any redundant node is not indispensable or dispensable. Therefore, the proposition holds.

### Pathway enrichment

Over-representation of protein in pathways was performed by using the Reactome Pathways database31. A p-value cut-off of 0.0001 and minimum matching proteins in pathway of 5 was used as the search parameters. Significant hits were ranked based on ration of protein matching, p-value, and False Discovery Rate (FDR), calculated using the Benjamin-Hochberg procedure36, which are adjusted p-values used to control the rate of false positives. Graphical representation of enriched pathways was obtained from Reactome database website.

## Acknowledgements

VR was supported by the Inspire Fellowship from the Department of Science and Technology (DST/INSPIRE/IF120809), the University of Manchester from a visiting researchers fund and is now supported by the Royal Society-SERB International Newton Fellowship (NF171560). VR, VS and GB thank DA-IICT for support, and GB also thanks IIIT-Delhi for support. DLR was partially supported by the MRC (MC_UU_1201412). DLR and JCN thank the Royal Society for an International Exchange grant (IE160248). JCN was partially supported by JSPS KAKENHI (JP25330351). TA was partially supported by JSPS KAKENHI (JP26540125). This research was also supported in part by Research Collaboration Projects of the Institute for Chemical Research, Kyoto University.

## Author contributions statement

V.R., J.C.N., J.-M.S., and D.L designed research; V.R., J.C.N., and A.O. performed research; T.A., M.I., V.S., G.B., and J.-M.S. contributed new analytical tools; V.R., J.C.N., T.A., M.I., and A.O. analysed data; V.R., J.C.N., and D.L. wrote the paper.

## Supplementary information

Supplementary Material: details of HCV analysis, and supplementary tables and figures.

Supplementary Note: details of theorems derived.

Supplementary Data file 1: network details.

Supplementary Data file 2: network analysis results.

Supplementary Data file 3: critical driver node comparison.

